# Depleting glioblastoma cells of very long-chain acyl-CoA synthetase 3 (ACSVL3) produces metabolic alterations in non-lipid pathways

**DOI:** 10.1101/2023.09.18.558236

**Authors:** Elizabeth A. Kolar, Xiaohai Shi, Emily M. Clay, Yanqiu Liu, Shuli Xia, Cissy Zhang, Anne Le, Paul A. Watkins

## Abstract

Knockout (KO) of the fatty acid-activation enzyme very long-chain acyl-CoA synthetase 3 (ACSVL3; SLC27A3) in U87MG glioblastoma cells reduced their malignant growth properties both in vitro and in xenografts. These U87-KO glioma cells grew at a slower rate, became adherence-dependent, and were less invasive than parental U87 cells. U87-KO cells produced fewer, slower-growing subcutaneous and intracranial tumors when implanted in NOD-SCID mice. Thus, depleting U87MG cells of ACSVL3 restored these cells to a phenotype more like that of normal astrocytes. To understand the mechanisms underlying these beneficial changes, we investigated several possibilities, including the effects of ACSVL3 depletion on carbohydrate metabolism. Proteomic and metabolomic profiling indicated that ACSVL3 KO produced changes in glucose and energy metabolism. Even though protein levels of glucose transporters GLUT1 and GLUT3 were reduced by KO, cellular uptake of labeled 2-deoxyglucose was unaffected. Glucose oxidation to CO_2_ was reduced nearly 7-fold by ACSVL3 depletion, and the cellular glucose level was 25% higher in KO cells. Glycolytic enzymes were upregulated by KO, but metabolic intermediates were essentially unchanged. Surprisingly, lactate production and the levels of lactate dehydrogenase isozymes LDHA and LDHB were elevated by ACSVL3 KO. The activity of the pentose phosphate pathway was found to be lower in KO cells. Citric acid cycle enzymes, electron transport chain complexes, and ATP synthase protein levels were all reduced by ACSVL3 depletion. Mitochondria were elongated in KO cells, but had a more punctate morphology in U87 cells. The mitochondrial potential was unaffected by lack of ACSVL3. We conclude that the beneficial effects of ACSVL3 depletion in human glioblastoma cells may result in part from alterations in diverse metabolic processes that are not directly related to role(s) of this enzyme in fatty acid and/or lipid metabolism. (Supported by NIH 5R01NS062043 and KKI institutional funds.)

## INTRODUCTION

Central nervous system (CNS) tumors represent the seventh most common cancer in adults in the United States, and about 80% of primary malignant CNS tumors are of glial origin (gliomas) (1). The World Health Organization (WHO) grades gliomas by pathologic evaluation, using a scale of I-IV based on cellular differentiation status (2). Glioblastoma multiforme (GBM; WHO grade IV) is the most commonly occurring malignant brain tumor, accounting for 14% of all tumors and 50% of all malignant tumors (1). Once diagnosed, prognosis for GBM is very poor. Despite standard-of-care treatment of GBM patients with surgery, radiation, and chemotherapy (temozolomide), the 5-year survival rate has remained at around 5% for many years (1). Therefore, new treatment strategies and new therapeutic targets are needed. The fatty acid metabolism enzyme ACSVL3 has emerged as a promising therapeutic target.

ACSVL3 (gene symbol SLC27A3) is an acyl-CoA synthetase (ACS) belonging to the “very long-chain” family; it is also known as fatty acid transport protein 3 (FATP3) (3,4). ACSs play a central role in lipid metabolism by forming a thioester between a fatty acid and coenzyme A (CoA) (5). The high-energy thioester bond facilitates metabolic reactions of the “activated” fatty acyl-CoA, allowing its participation in downstream anabolic (e.g. complex lipid synthesis) and catabolic (e.g. β-oxidation) pathways (5).

We found that ACSVL3 was highly overexpressed in human gliomas of all WHO grades, including GBM (6). Expression of ACSVL3 protein in the GBM-derived cell line U87MG was ∼10-fold higher than in primary human astrocytes [Watkins et al., unpublished observation]. To investigate the role of this protein in GBM malignancy, we knocked out ACSVL3 in U87MG cells, creating U87-KO cells (7). Compared to U87MG cells, U87-KO cells grew slower in culture and became adherence-dependent. Rapidly growing tumors were produced when U87MG cells were implanted subcutaneously in nude mice; in contrast, fewer xenografts developed from U87-KO cell implantation, and the tumors that did form grew at much slower rates (7). Thus, U87-KO cells display far less malignant behavior than do U87MG cells, both *in vitro* and *in vivo*.

To determine the mechanism(s) by which ACSVL3 supports the malignant phenotype of U87MG cells, we considered several hypotheses, including increased membrane phospholipid synthesis, changes in other lipid metabolic pathways, decreased apoptosis, alterations in autophagy, changes in cell cycle, and changes non-lipid metabolic pathways. We previously reported that while membrane phospholipid synthesis was not significantly different in U87-KO cells vs. U87MG cells, there were significant alterations in sphingolipid metabolism when ACSVL3 levels were depleted (7). We also reported that malignant U87MG cells were quite resistant to apoptosis, and that increased apoptosis in U87-KO cells could not explain the slower growth rate of these cells (8). In contrast, the cell cycle was dramatically altered in ACSVL3-deficient U87-KO cells (8), a phenomenon that may be related to the alterations in sphingolipid metabolism (7). In this report, we show that KO of ACSVL3 in U87MG cells produces significant changes in non-lipid pathways of intermediary metabolism as well.

## MATERIALS AND METHODS

### Materials and general methods

2-Deoxy-[^3^H]glucose was from PerkinElmer. D-[U-^14^C]glucose was from Moravek Biochemicals. Rabbit anti-Tom20 antibody was a gift from Dr. Hiromi Sesaki (Johns Hopkins Univ. Sch. Med.). Cy3 conjugated anti-rabbit secondary antibody was from Jackson Immunoresearch (211-1665-109). Protein was measured as described by Lowry et al. (9). The NAD^+^/NADH-Glo ™ Assay kit (Promega) was used following the manufacturer’s instructions. The MitoPotential kit (EMD Millipore) was used with the Muse Cell Analyzer following the manufacturer’s instructions.

### Cell lines and culture

The GBM cell line U87MG was from American Type Culture Collection (Manassas, VA). Authentication of this cell line was performed by the Johns Hopkins Genetics Resources Core Facility by short tandem repeat analysis using the PowerPlex 16 HS kit (Promega) within a year of study. Cell culture media was from Corning Cellgro, and fetal bovine serum (FBS) was from Gemini Bioproducts. Cells were cultured in MEM (minimum essential medium, Eagle) containing 10% FBS and supplemented with 10 mM HEPES, 1 mM pyruvate, and nonessential amino acids. Cells were incubated in a humidified incubator with a 5% CO_2_ atmosphere at 37°C. The ACSVL3-deficient U87-KO cell line was produced via the zinc finger nuclease strategy as previously described (7).

### Glucose uptake

Cells were cultured in duplicate 12-well plates until confluent. HEPES-buffered Krebs-Ringer solution (KRH) contained 0.5% bovine serum albumin, 25 mM HEPES, 120 mM NaCl, 5 mM KCl, 1.2 mM MgSO4, 1.3 mM CaCl2, and 1.3 mM KH2PO4. Labeling medium was prepared by adding 1 μl of 1 μCi/ml 2-deoxy-[^3^H]glucose per 1 ml KRH buffer and kept at 37°C. Cells were washed twice with warm Dulbecco’s phosphate-buffered saline (DPBS, containing calcium and magnesium). 1 ml of warm serum-free U87 culture medium was added to each well, and plates were returned to the CO_2_ incubator for 1 hour. Cells were washed twice with warm DPBS, and then incubated for an additional 20 min in 1 ml of warm KRH buffer. KRH buffer was aspirated, and cells were incubated in 0.5 ml of labeling media for 10 min at 37°C. Labeling buffer was aspirated, and the plates were put on a bed of ice to prevent further glucose uptake. Wells were washed twice with ice-cold DPBS after which cells were solubilized in 240 μl of warm 0.8% Triton by sonication (Branson 2150 ultrasonic cleaner) for 10 min at 37°C. Cell lysates were counted (150 μl) in a scintillation counter. Control 12-well plates were treated similarly, except that all procedures were done on ice. Specific activity of glucose uptake is reported as nmol/20 min/mg protein. Results are the mean ± SD for 5 separate determinations.

### Glucose oxidation to CO_2_

Cells were cultured in T-25 flasks until confluent. Labeling medium was prepared by adding D-[U-^14^C]glucose to pre-warmed KRH buffer for a final glucose concentration of 5 mM. On the day of the assay, media was aspirated from culture flasks and the cells were washed twice with warm (37°C) PBS. 5 ml of warm KRH buffer was added. After 20 min, the KRH buffer was aspirated from the flasks, and 5 ml of labeling medium was added. The flasks were capped with rubber stoppers fitted with hanging wells containing Whatman glass microfiber filter paper that had been wetted with 20 ml of 1 M KOH to trap the ^14^CO_2_ produced. The flasks were incubated for 1 hour at 37°C with gentle shaking. Glucose metabolism was terminated by injection of 0.25 ml 60% perchloric acid through the rubber stopper, and the flasks were kept at 4°C overnight with gentle shaking to trap the ^14^CO_2_. The filter paper was transferred to scintillation vials and 10 ml of scintillation cocktail was added. Radioactivity in the vials was quantitated by scintillation counting, and specific activity (nmol/20 min/mg protein) was calculated. Results are the mean ± SD for 4 separate determinations.

### Immunofluorescence

Immunofluorescence was performed by culturing U87MG and U87-KO cells on glass coverslips until nearly confluent. The coverslips were then washed twice with 1x PBS and fixed with 4% formaldehyde in PBS for 20 min at room temperature. The fixation buffer was aspirated, and the coverslips were washed three times with PBS (5 min each wash). Cells were permeabilized with 0.1% Triton X-100 in PBS for 8 min at room temperature, after which the coverslips were washed with PBS as above. Anti-Tom20 antibody was diluted to a final concentration of 1:400 in 0.5% BSA in 1x PBS. 20 μl of antibody was pipetted onto parafilm and the coverslips were gently placed upside down on the antibody dilution. Coverslips were incubated overnight at 4°C in a humidified chamber. Coverslips were washed as above and incubated with anti-rabbit secondary antibody conjugated to Cy3 (1:250) for 1 hour at room temperature. After washing three times with PBS, coverslips were mounted onto slides. Fluorescent images were captured using a Zeiss AxioImager fluorescent microscope with ApoTome attachment, and analyzed using Adobe PhotoShop CS5.

### Lactate secretion

Cells were cultured to near confluence in 5 cm dishes. At times 0, 0.5, 1, 2, 4, and 6 hours, aliquots of culture medium were removed and kept on ice, Samples were assayed for lactate concentration essentially using the method originally described by Gutmann and Wahlefeld (10). Cuvettes contained (final concentrations, in a total volume of 1 ml) hydrazine/glycine (0.18 M/0.45 M), pH 9.5, NAD (2 mg/ml), and sample. The absorbance at 340 nm was measured (baseline), and lactate dehydrogenase (0.1 mg, Sigma) was added. After 10-20 min at room temperature, the increase in absorbance was measured. The extinction coefficient for NAD (6.22 cm^2^/μmol) was used to calculate the concentration of NAD in the medium. Results are presented as mean of triplicate assays ± SD.

### G6PD assay

To estimate glucose metabolism via the pentose phosphate pathway, the activity of the initial, rate-limiting enzyme, glucose-6-phosphate dehydrogenase, was assayed. The assay originally described by Löhr and Waller (11) was adapted for use in a 96-well plate. Cells were solubilized in 0.5% Tween-20. Reactions contained (final concentrations) 50 mM triethanolamine, pH 7.5, 5 mM disodium EDTA (in 1% NaHCO3), 0.5 mM MgCl2, 0.5 mM NADP^+^, and solubilized cells (0.02-0.04 mg protein). The reaction was started by addition of glucose-6-phosphate (final concentration, 0.7 mM), and appearance of NADPH was monitored as fluorescence increase using a Spectra Max plate reader (λex = 340 nm; λem = 450nm; λcutoff = 420nm). Results are expressed as relative fluorescence units (RFU) per min per mg protein.

### Mitochondrial potential

The Muse Cell Analyzer was used with the MitoPotential kit following the manufacturer’s instructions. This assay measures the percentage of a cell population that is dead, depolarized, or both based on the integrity of the mitochondrial membrane.

### Metabolomic analysis

Cells were seeded onto 150 mm culture dishes and grown to 80-90% confluence. After washing twice with ice-cold PBS, dishes were placed on dry ice and cells were scraped into a total of 1.05 ml of MeOH/water (2.5:1) and transferred to a 2 ml microcentrifuge tube. Cells were sonicated by five one-second pulses on ice, followed by the addition of 0.375 ml chloroform and mixing. 0.375 ml water was added and tubes were vortexed. After addition of 0.175 ml chloroform and mixing, tubes were centrifuged for 30 min at 3,000 x g. After centrifugation, the samples separate into 3 layers. Each layer was separately dried using a SpeedVac. The top layer (polar compounds) was analyzed using Q-TOF LC/MS with data analysis performed with MassHunter Qualitative Analysis and Mass Profiler Professional software. Protein concentration was determined on the middle layer. The bottom layer (lipids) was saved for future lipidomic studies. Results are presented as mean fold change in U87-KO cells with respect to the U87MG group +/- SEM for triplicate analyses of duplicate dishes. Data were normalized using the protein concentration.

### Proteomic analysis

Proteomic profiling was performed exactly as previously described (7,12,13). Tryptic peptides were labeled with tandem mass tags, fractionated by reverse-phase liquid chromatography, and fractions analyzed using an Orbitrap Elite tandem mass spectrometer. Data from biological replicates was obtained and the fold change in U87-KO vs. U87MG was calculated.

### Statistics

Statistical significance was determined using Student’s t-test. All results are presented as mean ± SD for the number of observations specified, except for metabolomic data where mean ± SEM is shown.

## RESULTS

### Glycolytic flux is paradoxically increased in ACSVL3-deficient U87-KO cells

For energy production, many tumors rely on glycolysis rather than the complete oxidation of glucose in the mitochondrion despite adequate availability of oxygen (“aerobic glycolysis”), a condition first described by Otto Warburg nearly a century ago (14). Thus, we suspected that depleting U87MG cells of ACSVL3 - which produces a more normal growth phenotype - would render them more oxidative and less dependent on glycolysis. To address this question, we used a combination of metabolomic, proteomic, and enzyme/pathway assay approaches.

Analysis of intracellular analytes indicated that the glucose level in U87-KO cells was higher than that of U87MG cells (Fig. 1A; *p*=0.009). Proteomics detected two glucose transporters, GLUT1 and GLUT3, in these cells (Table 1); levels of both transporters were decreased in cells lacking ACSVL3. However, despite decreased transporter levels, the rate of glucose uptake measured using 2-deoxy-[^3^H]glucose was identical in both U87MG and ACSVL3-deficient U87-KO cells (Fig. 2). To resolve this discrepancy, we assessed flux through glycolysis by measuring the rate of lactate secretion into the cell culture medium. Surprisingly, lactate secretion was higher in KO cells than in U87MG cells, suggesting that glycolytic flux was higher, rather than lower, when ACSLV3 was depleted (Fig. 3). Proteomic analysis supported this conclusion, as levels of both lactate dehydrogenase subunits, LDHA and LDHB, were elevated in KO cells (Table 1). Intracellular lactate levels trended higher in U87-KO cells, but differences did not reach statistical significance (data not shown). Intracellular pyruvate levels were not affected by KO of ACSVL3 (data not shown).

**Table 1.**
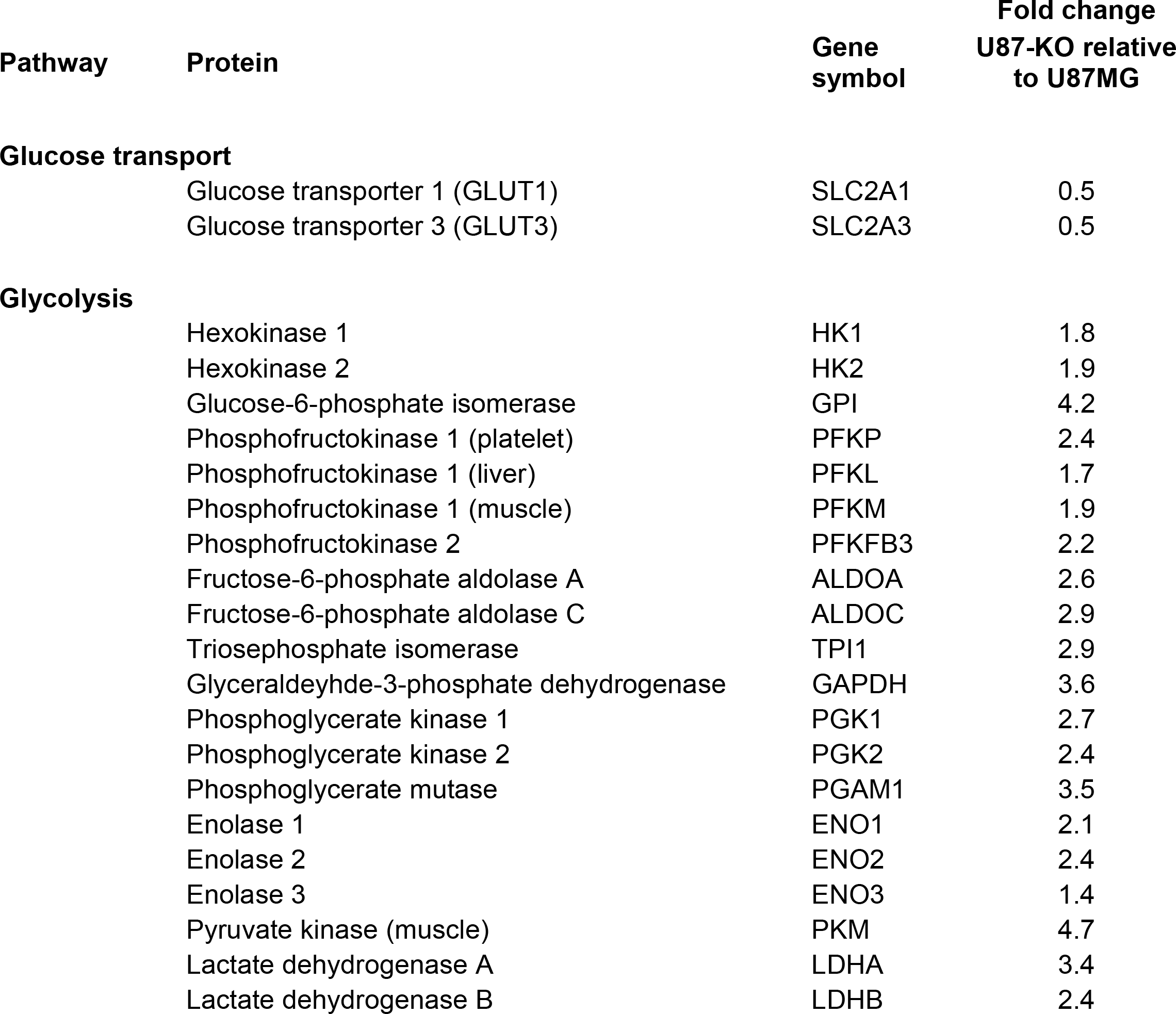
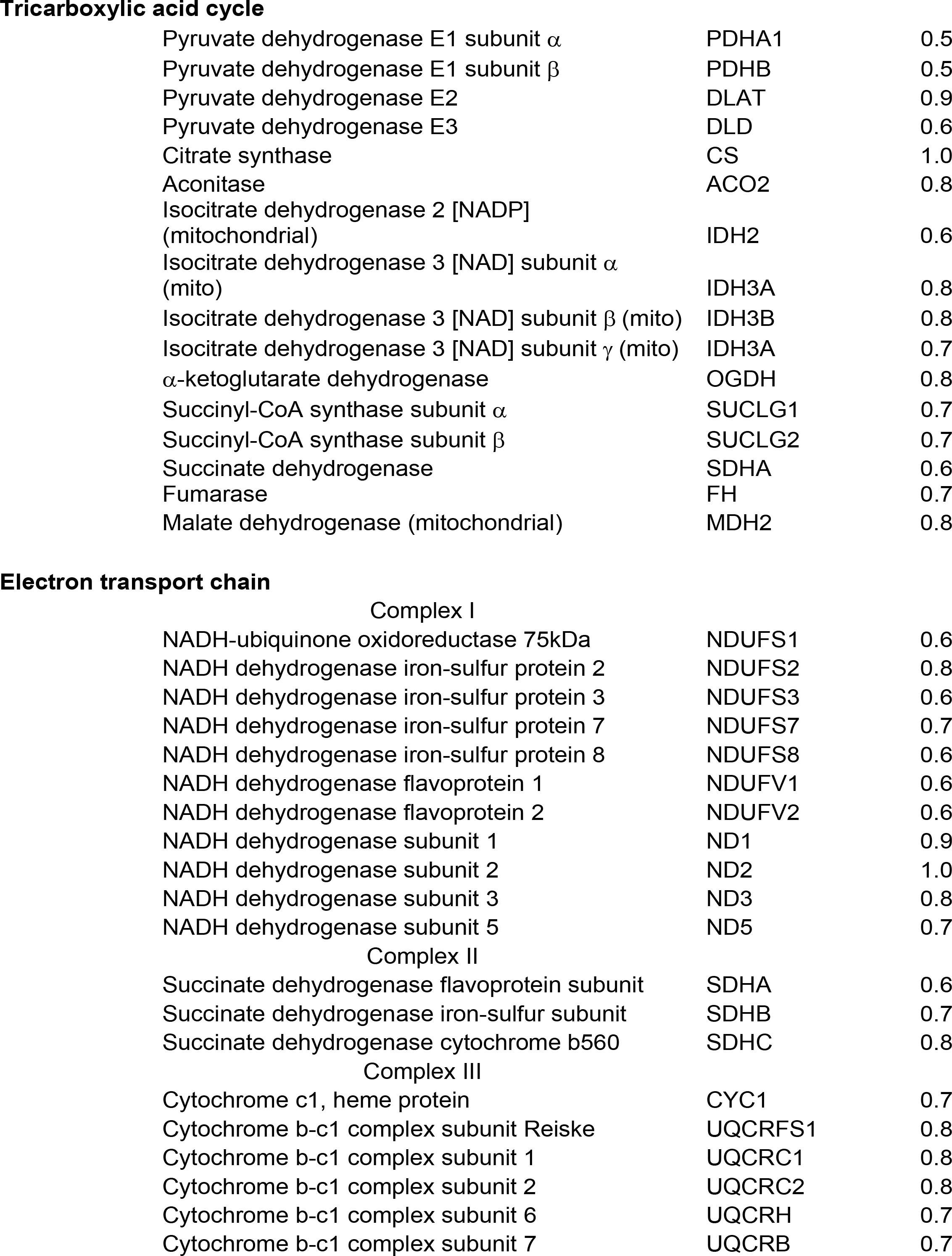
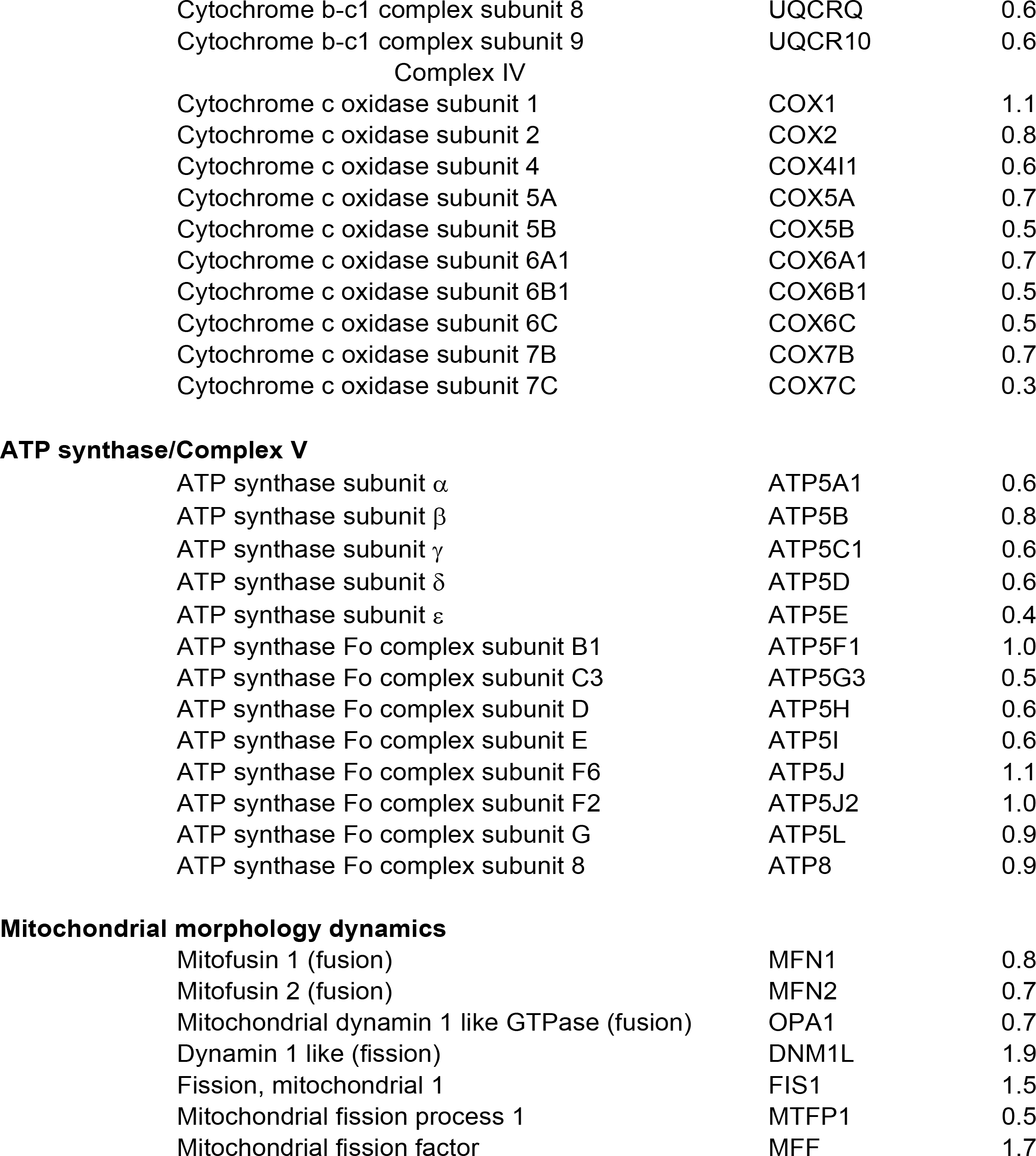
Proteomic analysis of U87MG and U87-KO cells. The relative abundance of proteins relevant to glucose metabolism, TCA cycle, OXPHOS, and mitochondrial dynamics in U87-KO relative to U87MG cells was determined by tandem mass spectrometry as described in Methods.

**Figure 1.**
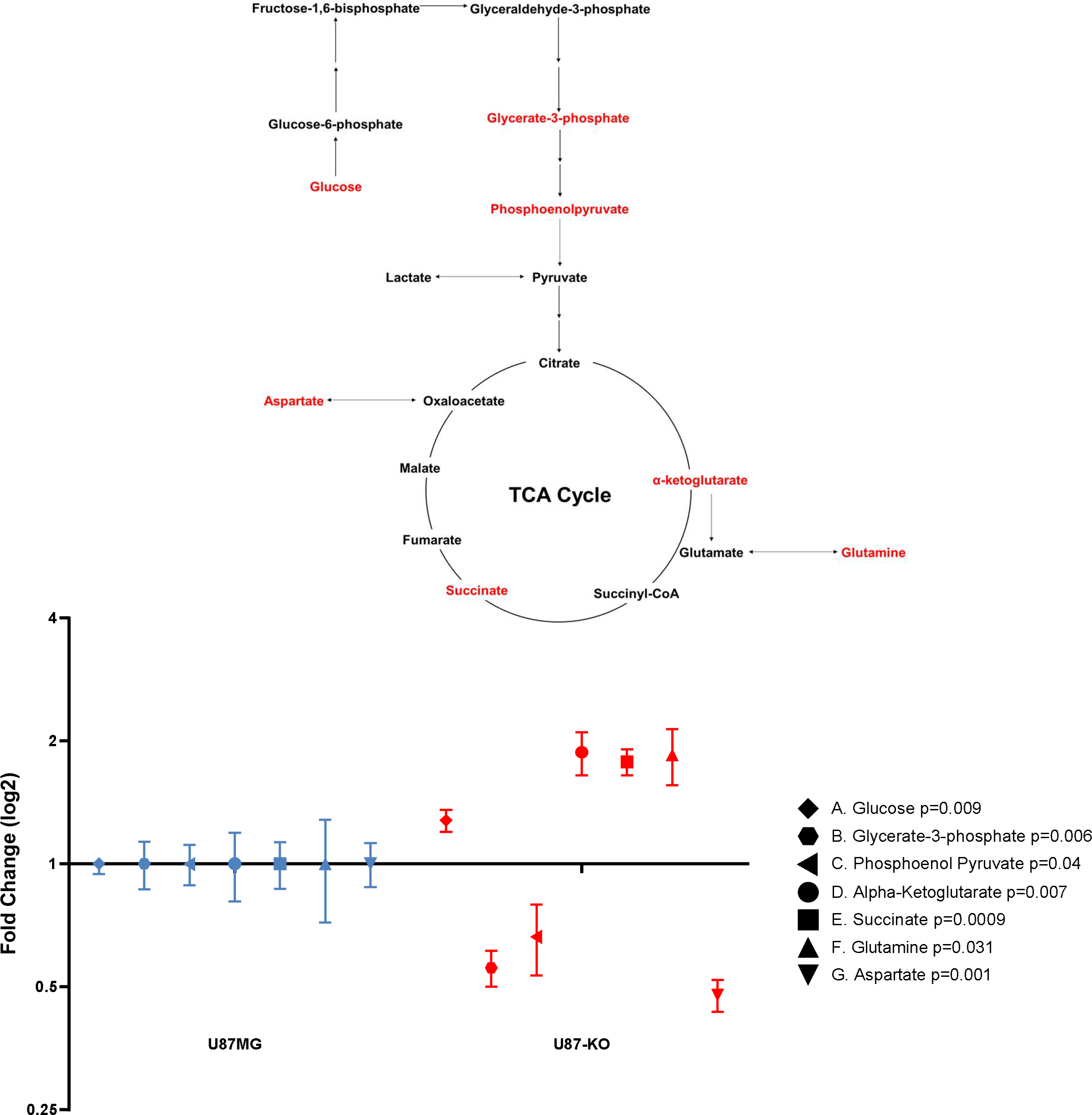
Metabolomic analysis of U87MG vs. U87-KO cells. Cells were grown to near confluence and processed for analysis of metabolites by Q-TOF LC/MS as described in Methods. Top, glycolytic and TCA cycle pathways. Only those intermediates detected by metabolomic analysis are shown. Metabolites whose levels were significantly altered by ACSVL3 KO are in red typeface. Bottom, mean fold change of metabolite levels in U87-KO with respect to U87MG cells ± SEM for triplicate analyses of duplicate dishes of cells. Metabolites with significantly altered levels included three glycolytic intermediates (A-C), two TCA intermediates (D, E), and two related intermediates (F, G) are shown. Blue, U87MG; red, U87-KO.

**Figure 2.**
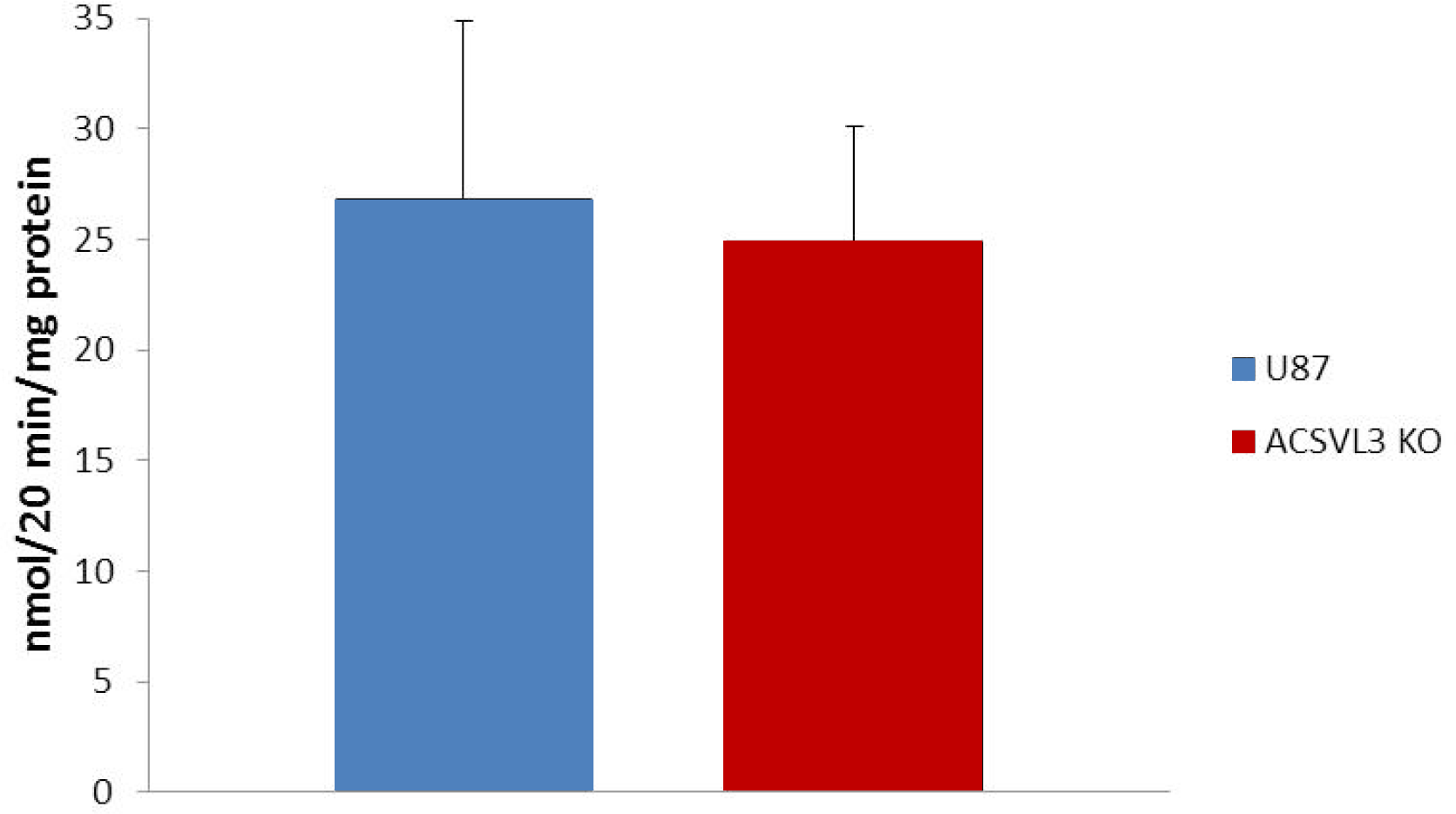
Glucose uptake in U87-KO cells relative to U87MG cells. Cells were incubated in medium that included 2-Deoxy-[^3^H]glucose and the radioactivity in aliquots of culture medium was determined. Results are presented as nmol glucose uptake/20 min/mg protein (mean ± SD; n = 5). Blue bars, U87MG; red bars, U87-KO.

**Figure 3.**
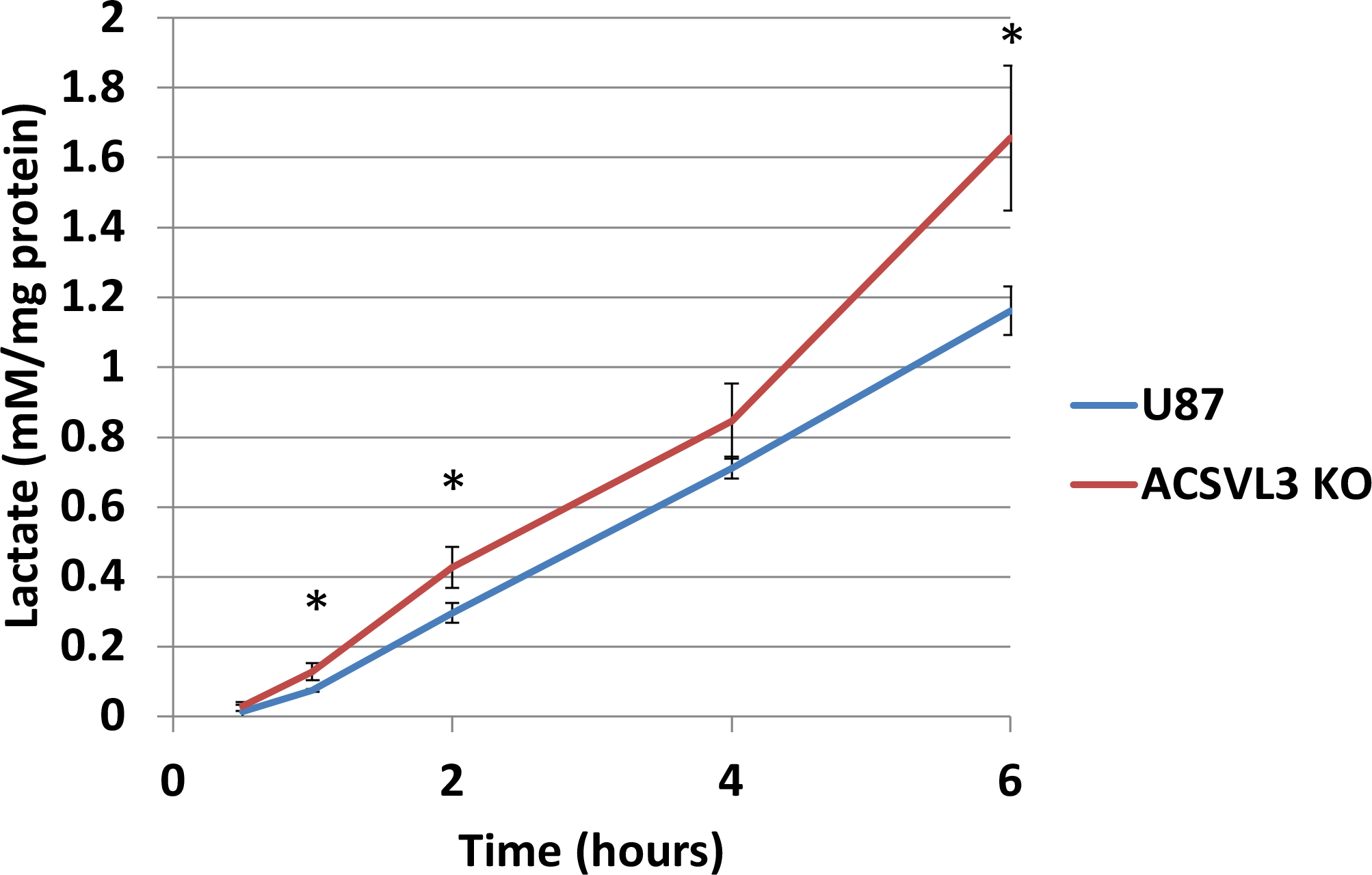
Lactate secretion into culture medium. Cells were cultured to near confluence. The culture medium was replaced and aliquots were removed at the indicated times for assay of lactate concentration. Results are presented as mM/mg protein (mean ± SD) for triplicate assays. Blue lines, U87MG; red lines, U87-KO. Asterisks indicate statistical significance at the level of *p*<0.05.

In support of the conclusion that glycolysis is increased by depletion of ACSVL3, levels of most glycolytic enzymes were found to be higher in ACSVL3 KO cells, including hexokinases 1 and 2, glucose-6-phosphate isomerase, phosphofructokinase 1, phosphofructokinase 2, fructose-6-phosphate aldolases A and C, triosephosphate isomerase, glyceraldeyhde-3-phosphate dehydrogenase, phosphoglycerate kinases 1 and 2, phosphoglycerate mutase, enolases 1-3, and pyruvate kinase (muscle isozyme) (Table 1).

Similarly, levels of glycolytic intermediates (Fig. 1) were either unchanged or lower when U87MG cells lacked ACSVL3, as would be expected if flux through the pathway was increased. Out of the glycolysis intermediates detected, levels of glycerate-3-phosphate (Fig 1B; *p*=0.006) and phosphoenolpyruvate (PEP) (Fig. 1C, p=0.04) were lower in U87-KO cells compared to U87MG cells; differences between levels of other intermediates did not reach statistical significance.

### Glucose oxidation to CO_2_ is reduced when ACSVL3 is depleted in U87 cells

Under aerobic conditions, complete degradation of glucose requires metabolic processes in the mitochondrion including tricarboxylic acid (TCA) cycle, electron transport chain and oxidative phosphorylation (OXPHOS), where oxidation of the end product of glycolysis, pyruvate, to CO_2_ and water is coupled to ATP production. To assess the overall metabolic profile, we measured the conversion of [U-^14^C]glucose to ^14^CO_2_, which requires both glycolysis and the activity of the TCA cycle. In contrast to our original prediction that oxidative metabolism of glucose would be higher in ACSVL3-depleted cells, the rate of CO_2_ production was significantly lower in KO cells (Fig. 4). Correspondingly, levels of most TCA cycle enzymes were decreased by ACSVL3 KO (Table 1); only citrate synthase levels were unchanged by depletion of ACSVL3. Metabolomic analysis indicated a potential bottleneck in the TCA cycle at the level of succinate dehydrogenase (SDH) (Fig. 1). Levels of TCA cycle intermediates prior to SDH including α-ketoglutarate (Fig. 1D; *p*=0.007) and succinate (Fig. 1E; *p*=0.0009) were significantly elevated in KO cells. However, downstream of SDH, no TCA cycle intermediate level was statistically different between the 2 groups. In addition, glutamine was also significantly higher in the U87-KO cells (Fig. 1F; *p*=0.031); transamination of the TCA intermediate α-ketoglutarate yields glutamate, which is amidated to produce glutamine. On the contrary, aspartate, a product of oxaloacetate transamination, was significantly lower in the U87-KO cells (Fig. 1G; *p*=0.001). This helped further confirm the trend observed in the TCA cycle intermediates.

**Figure 4.**
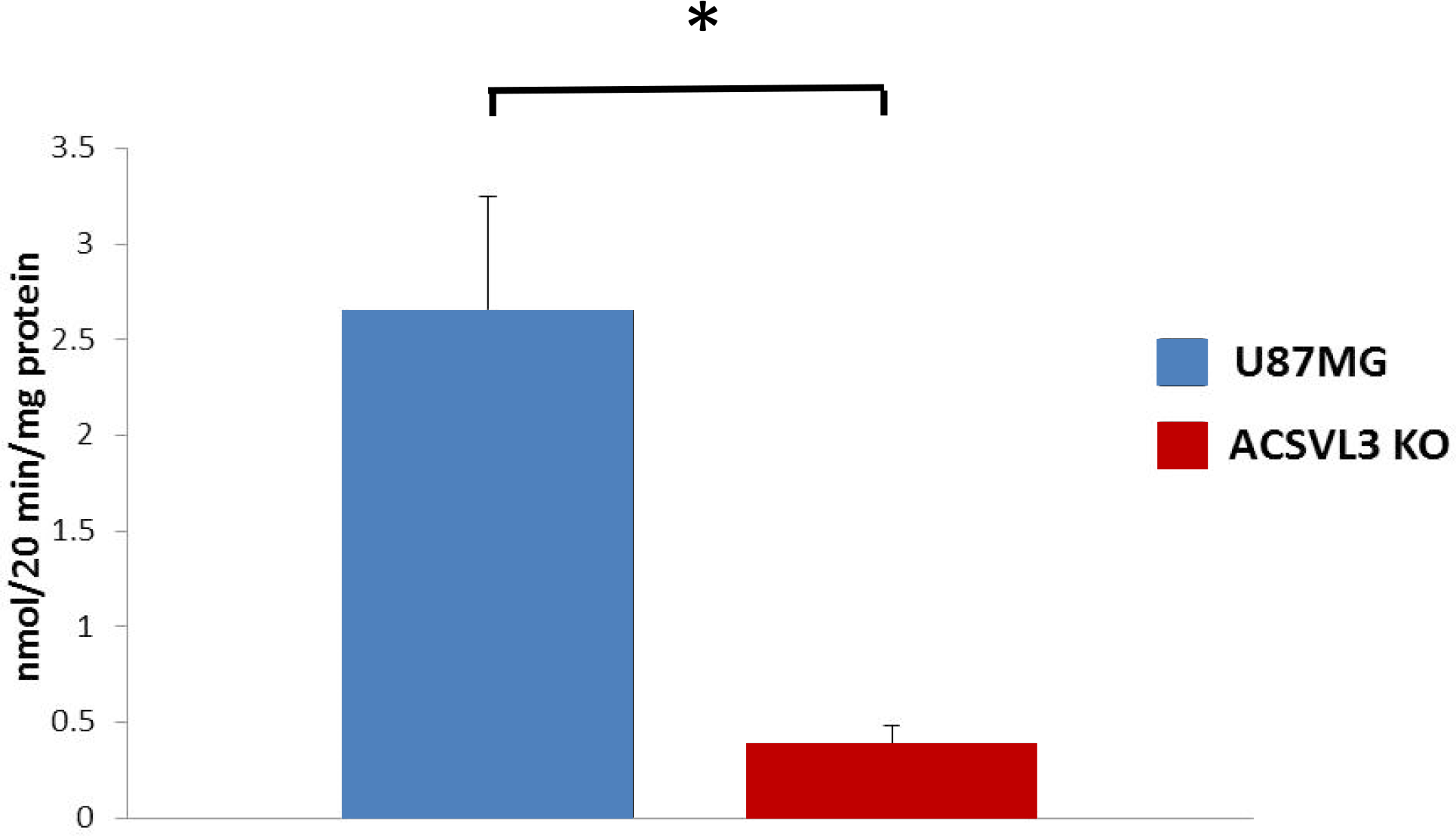
Oxidation of glucose in U87MG and U87-KO cells. Cells of both lines were incubated with uniformly labeled D-[14C]glucose and release of labeled CO_2_ was quantitated. Results are presented as nmol 14CO_2_ produced/20 min/mg protein (mean ± SD; n=4). Blue bars, U87MG; red bars, U87-KO. The difference was significant at the level *p* = 0.006.

Proteomics also revealed that KO of ACSVL3 in U87MG cells was associated with decreased levels of many proteins required for the electron transport chain and OXPHOS (Table 1). Of 32 constituent proteins in complexes I, II, III, and IV of the respiratory chain, levels of all but 2 were lower in U87-KO cells. Five subunits of ATP synthase F1 complex were all lower in U87-KO cells. Although the levels of 5 subunits of ATP synthase FO complex were also decreased by lack of ACSVL3, two others were unchanged and another was slightly elevated.

### Pentose phosphate pathway activity is lower in U87-KO cells

In addition to degradation via glycolysis, glucose can also be metabolized via the pentose phosphate pathway (PPP). The PPP provides substrates for nucleotide synthesis and reducing equivalents for biosynthetic processes, both of which are important in cancer cell metabolism. To assess PPP function, we measured the enzyme activity of glucose-6-phosphate dehydrogenase (G6PD), the first and rate-limiting step of the pathway. As shown in Fig. 5, G6PD activity was significantly lower in ACSVL3-deficient U87-KO cells (*p* < 0.05) than in U87MG cells, suggesting decreased flux through PPP. This observation is consistent with the reduced growth rate of U87-KO cells.

**Figure 5.**
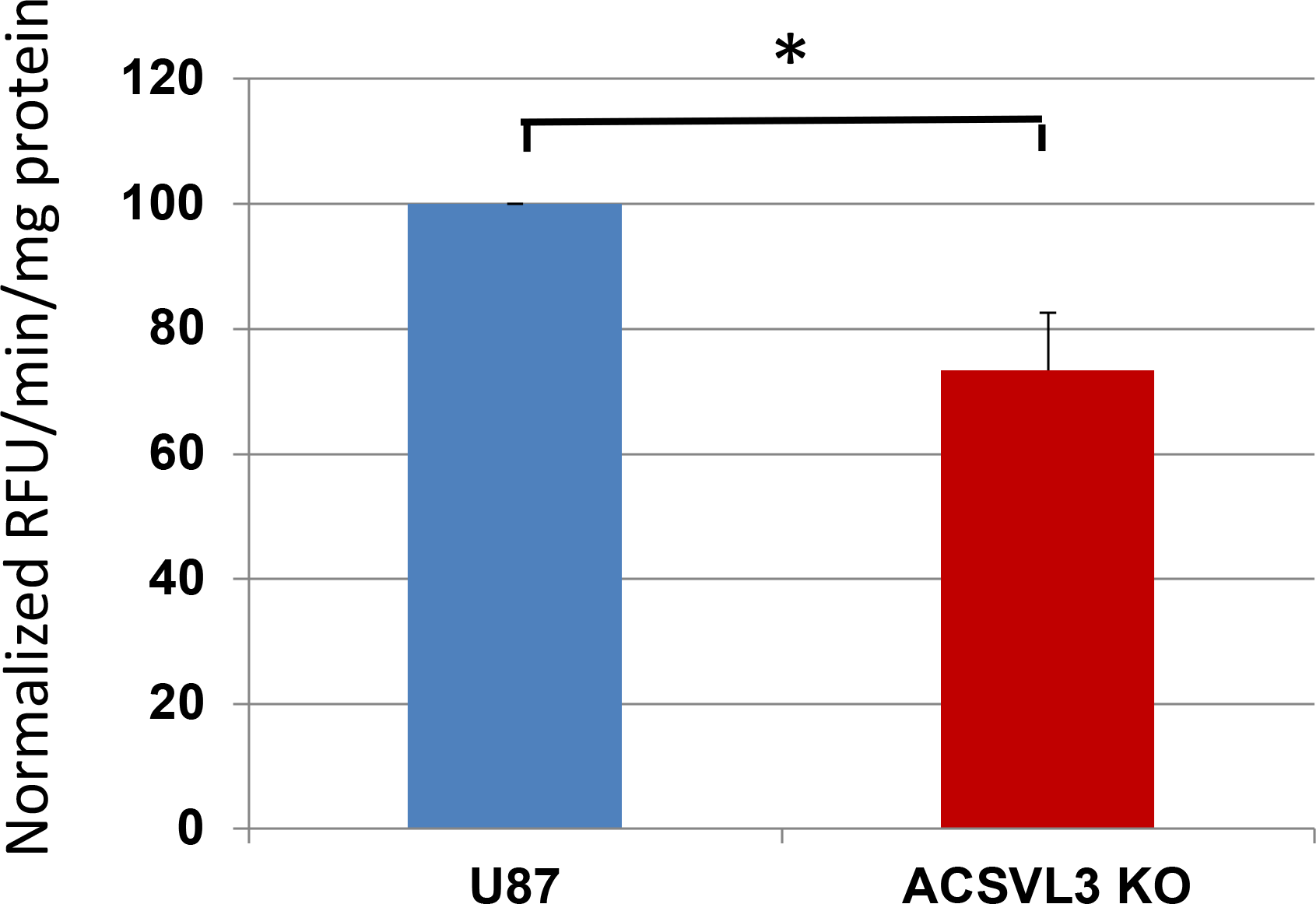
Glucose-6-phosphate dehydrogenase enzyme activity in U87MG and U87-KO cells. G6PD activity was determined as a surrogate for pentose phosphate pathway activity. Measurement of G6PD-dependent conversion of NADP+ to NADPH was quantitated fluorimetrically. Results are expressed as relative fluorescence units (RFU) per min per mg protein, normalized to U87MG cells (mean ± SD; n=3). Blue bars, U87MG; red bars, U87-KO. The difference was significant at the level *p* < 0.05.

### Mitochondrial function and morphology are altered by ACSVL3 depletion in U87 cells

The above studies of glucose homeostasis and energy metabolism suggested that depletion of ACSVL3 could affect mitochondrial function and/or morphology. Energy metabolism is regulated to a great extent by redox reactions involving NAD^+^ and NADH. Up to 70% of the cellular NAD^+^ pool is thought to be mitochondrial (15-17). Therefore, we measured the NAD^+^/NADH ratio in U87MG cells and ACSVL3 KO cells and found it to be significantly elevated (*p* < 0.05) when ACSVL3 was depleted (Fig. 6). A high NAD^+^/NADH ratio is also consistent with reduced activity of the TCA cycle, and reduced OXPHOS capacity.

**Figure 6.**
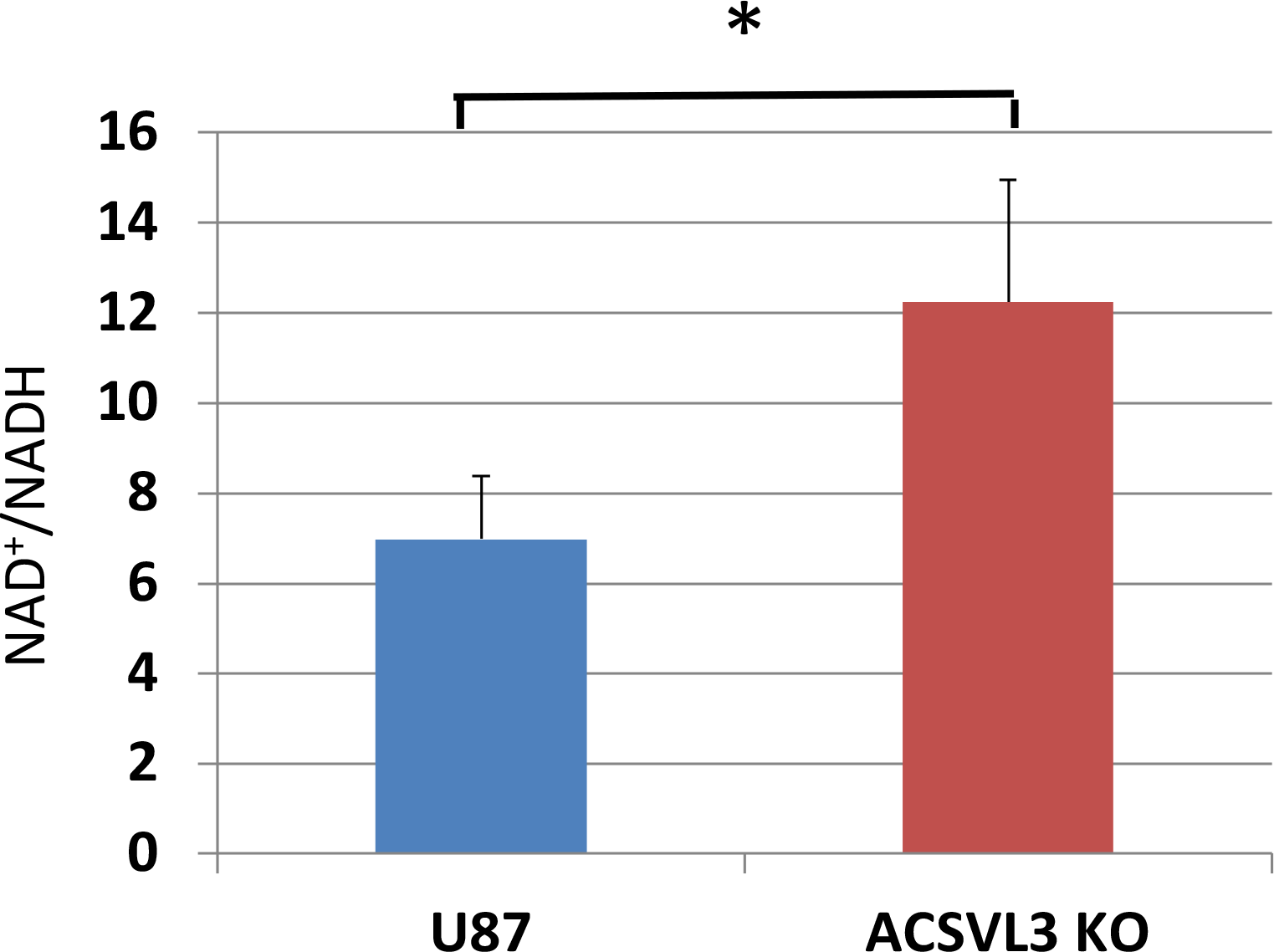
NAD^+^/NADH ratio in U87MG and U87-KO cells. The redox state of U87MG vs. U87-KO cells was measured using the NAD^+^/NADH-Glo™ Assay kit (Promega). Results are the mean ± SD for n=3 determinations. Blue bars, U87MG; red bars, U87-KO. The difference was significant at the level *p* < 0.05.

Lack of ACSVL3 could also be affecting U87MG cell growth by opening the mitochondrial transition pore. This would lead to depolarization of the mitochondrial membrane. Therefore, we measured mitochondrial membrane potential using a Muse Cell Analyzer. As shown in Fig. 7, the majority (96-97%) of both U87MG and U87-KO cells were alive, and of these only 1% of each were depolarized. These data indicate that both cell types contain functionally intact mitochondria.

**Figure 7.**
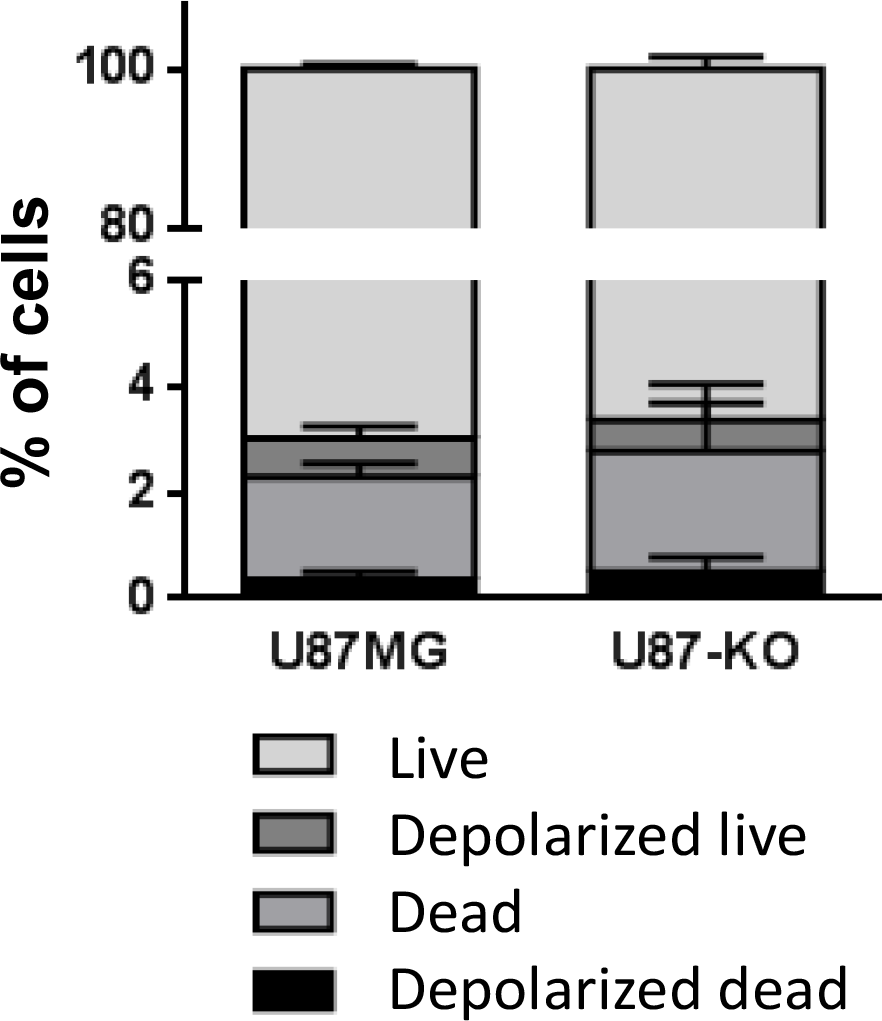
Mitochondrial membrane potential in U87MG and U87-KO cells. Mitochondrial membrane potential was measured by a flow cytometric method using the Muse® cell analyzer and MitoPotential kit. The assay quantitates the percentage of cells that are alive, alive but depolarized, dead, and depolarized dead. Results shown are mean ± SD for three determinations.

Mitochondrial morphology is a dynamic process that is related to metabolic state and bioenergetics; thus, changes in mitochondrial structure are often seen in cancer cells (18). Mitochondrial morphology was assessed by immunofluorescence analysis using a rabbit antibody against Tom20, a mitochondrial outer membrane transporter. We observed striking differences in mitochondrial morphology between U87MG and U87-KO cell lines. The mitochondria in the U87MG cell line appear more fragmented, while the mitochondria in the ACSVL3-deficient U87-KO cells are more fused and form reticular structures (Fig. 8). Changes in mitochondrial structure are regulated by proteins involved in fusion and fission of the organelle, and several of these were detected by proteomic analysis. Although decreases in proteins involved in fusion were detected in ACSVL3 KO cells, there was a more profound decrease in a critical fission protein, Mtfp1, that may account for the observed differences in morphology (Table 1).

**Figure 8.**
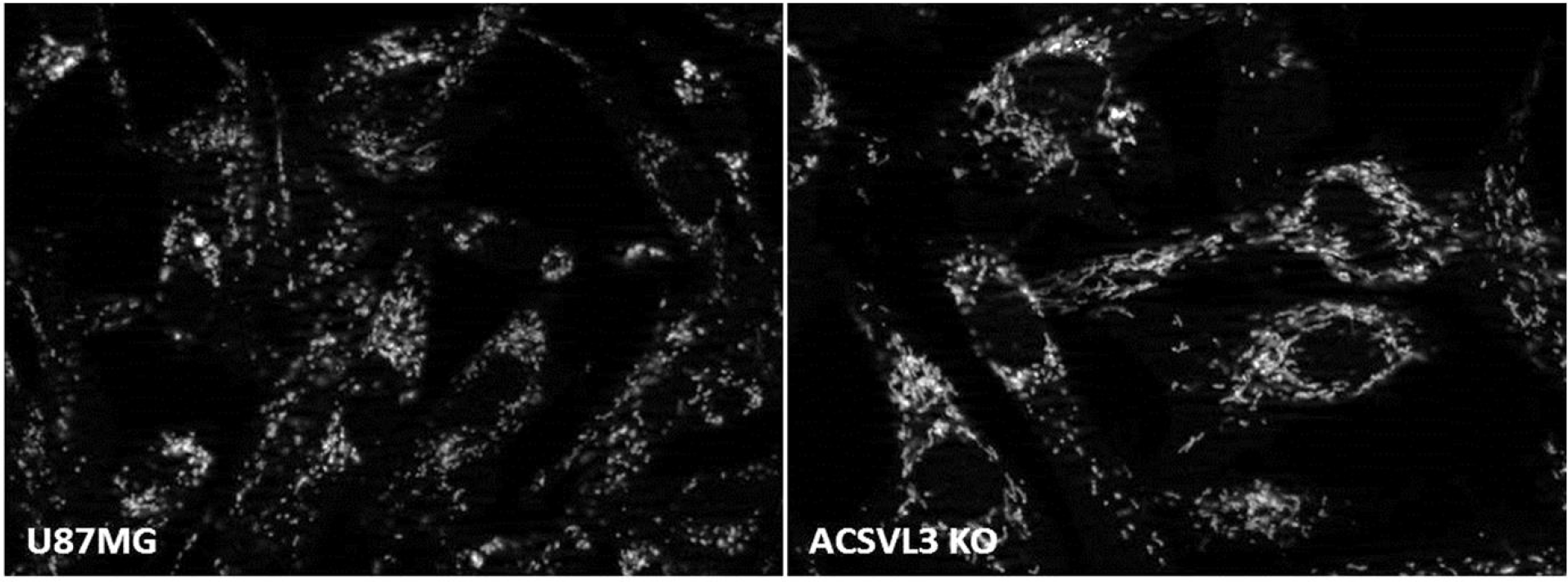
Immunofluorescence of TOM20 in U87MG and U87-KO cells. Mitochondrial morphology was assessed by fluorescence microscopy using an antibody against the outer membrane import receptor protein, TOM20, as described in Methods. U87MG cell mitochondria appear more punctate while the mitochondria of the U87-KO cells are more filamentous.

In summary, depletion of the fatty acid activating enzyme ACSVL3 in glioblastoma cell line U87MG, which results in a more non-malignant cellular phenotype, causes significant changes not only in lipid metabolism (7), but also in non-lipid pathways such as carbohydrate and energy metabolism. Further investigation will be necessary to understand fully the mechanistic underpinnings of these observations.

## DISCUSSION

Previous studies have clearly demonstrated that the fatty acid metabolism enzyme, ACSVL3, is overproduced in human gliomas, and in U87MG glioma cells (6,7). In addition to a rapid growth rate, the upregulated expression of ACSVL3 in U87MG cells was associated with alterations in fatty acid and lipid metabolism, as expected (7). In particular, ACSVL3 overexpression resulted in increased sphingolipid synthesis. We hypothesized that this contributes to the rapid growth phenotype of these glioblastoma cells (8). Depletion of ACSVL3 in U87 cells restores them to a more normal, less malignant growth phenotype, suggesting that targeting of this enzyme may have therapeutic benefits.

While changes in U87MG cell lipid metabolism were expected when ACSVL3 was depleted by genetic knockout, effects on other metabolic pathways, e.g. carbohydrate metabolism, were somewhat unanticipated. Regulatory links between glucose and lipid metabolism are important control mechanisms for overall cellular energy homeostasis. For example, the TCA cycle is an important crossroad between glucose and lipid metabolism. Pyruvate, the end-product of glycolysis, is metabolized by the mitochondrial pyruvate dehydrogenase complex and then enters the TCA cycle as acetyl-CoA. Acetyl-CoA is also produced when fatty acids are degraded by mitochondrial β-oxidation. Acetyl-CoA from either pathway can be completely oxidized to CO_2_ for energy production, or can be used as building blocks for fatty acid and cholesterol synthesis. More complex regulatory mechanisms linking glucose and lipid metabolism are also known. For example, Cheng et al. showed that glucose can regulate lipid metabolism through N-glycosylation of SCAP (SREBP-cleavage activating protein) to activate transcription factor SREBP-1 that induces lipid metabolism genes such as fatty acid synthase and the LDL receptor (19). Aberrant glucose metabolism in cancer has been an extensive subject of study since Otto Warburg made his observation that cancer cells have increased glycolysis, even in the presence of oxygen (14,20). Findings presented here show that ACSVL3 KO in the U87MG cell line affects glucose metabolism and mitochondrial structure and function.

Because of the typical observation of increased glycolysis in cancer cells, our working hypothesis was that ACSVL3 depletion would reduce glycolysis and increase glucose flux through TCA and OXPHOS. Indeed, Wang et al. reduced U87MG growth rate by targeting insulin-like growth factor receptor, and reported that both glycolysis and glycolytic capacity were decreased (21). Similarly, when Zhang et al. targeted the transcription factor E2F2 and slowed U87MG growth, both glucose consumption and lactate production were reduced (22). In contrast, reducing the growth rate by KO of ACSVL3 in U87 cells increased glycolysis, as measured by lactate accretion in the culture medium. This suggests that the mechanism of ACSVL3 reduction on U87MG growth is significantly different than expected. A study by Beckner et al. found that U87MG cells can utilize lactate, the end product of glycolysis, for gluconeogenesis and glycogen synthesis (23). It is possible that depleting cells of ACSVL3 somehow blocks flux of lactate through one or both of these pathways, and shunts more lactate into the culture medium.

In addition to increased lactate secretion by U87-KO cells, we found decreased activity of the TCA cycle and OXPHOS. Pyruvate dehydrogenase (PDH) is a key regulator of mitochondrial pyruvate metabolism. Phosphorylation of PDH by PDH kinase 1 (PDHK1) inactivates the PDH complex, thereby decreasing flux through the TCA cycle (24). Phosphoglycerate kinase 1 (PGK1), the first ATP-generating enzyme in the glycolytic pathway, can translocate from cytoplasm to mitochondria and act as a protein kinase to phosphorylate and activate PDHK1 (25). This phosphorylation would inhibit further pyruvate metabolism and enhance lactate production. Others have shown that PGK1 is upregulated in several cancers, including radioresistant astrocytomas (26). Interestingly, in this report we show that PGK1 was 2.7-fold higher in U87-KO cells (Table 1). This could suppress TCA cycle-mediated glucose/pyruvate oxidation even further, in agreement with results shown in Fig. 4, and increase lactate production.

The increase in the level of succinate when ACSVL3 expression was reduced was unexpected. Succinate has been reported to be an oncogenic metabolite that promotes epithelial/mesenchymal transition, invasion, and angiogenesis [27]. Decreasing, not increasing, succinate levels would be more consistent with the less malignant properties of U87-KO cells. Levels of succinate dehydrogenase A (SDHA) were, however, lower in U87-KO cells, consistent with elevated succinate.

The decrease in PPP activity in U87mg cells when ACSVL3 is depleted is consistent with decreased cell proliferation. PPP activity yields the ribonucleotide precursors of RNA and DNA required for rapid cell division exhibited by cancer cells. To function, the PPP requires cytoplasmic NADP^+^. When active lipogenesis (fatty acid synthesis) and cholesterogenesis is occurring, NADPH is consumed and NADP^+^ is generated. Decreased glucose metabolism via the PPP suggest a possible slowdown of lipid synthesis when ACSVL3 is depleted. While we found no major reduction in fatty acid and phospholipid synthesis from the 2-carbon precursor, acetate, in U87-KO cells (7), cholesterol synthesis was lower in these cells [EA Kolar, unpublished observation]. Whether this is sufficient to reduce availability of cytoplasmic NADP^+^ remains to be determined.

Depletion of ACSVL3 resulted in elevation of the NAD^+^/NADH ratio in U87 cells. Since mitochondria contain about 70% of the cellular NAD^+^ pool (15-17), it is possible that changes in mitochondrial NAD^+^ may affect glucose homeostasis. Van Linden et al. reported that artificially elevating mitochondrial NAD^+^ levels in HEK293 cells resulted in dramatic growth retardation and a metabolic shift from oxidative phosphorylation to glycolysis (28). The increased NAD^+^/NADH ratio is consistent with the increase in glycolysis and decrease in glucose oxidation observed in U87-KO cells. Although the mechanism underlying the increased NAD^+^/NADH ratio has not been identified, inhibition of PDH by phosphorylation as described above is consistent with this shift toward a more oxidized state for this redox nucleotide.

We also report here that KO of ACSVL3 altered the appearance of mitochondria. We previously reported that endogenous ACSVL3 partially colocalized with mitochondria in MA-10 and Neuro2a cells (29). U87-KO cells have network-like mitochondria, whereas U87MG cells have more punctate mitochondria. Mitochondrial morphology, particularly the dynamics of the mitochondria, has been shown to play an important roles in cancer metabolism, growth, and metastasis. While mitochondria in normal cells generally have an elongated, reticular appearance, tumor cell mitochondria are typically fragmented (30). It has been proposed that such changes in mitochondrial shape may be a key mechanism for promoting shifts in glucose, lipid, and amino acid metabolism in tumor cells (30). Our observations in U87MG and U87-KO cells are consistent with a more normal appearance when ACSVL3 levels are reduced.

Mitochondrial morphology is controlled by fission and fusion of the organelle, and these processes can be dysregulated in cancer. One study showed that human lung cancer cell lines exhibited excess mitochondrial fission and impaired mitochondrial fusion (31). An examination of the proteomic data revealed that, contrary to our expectations, fusion proteins are low in the U87-KO cells, but fission proteins are higher, except for Mtfp1, also known as Mtp18. This protein was found to be essential for mitochondrial fission, as RNAi-mediated loss of function resulted in highly fused mitochondria, even with an overexpression of another fission protein, Fis1 (32). This decrease in Mtp18 may be sufficient to prevent fission and keep mitochondria in a highly fused form, even when other fission proteins are overexpressed. To address whether changes in structure affected the integrity of the organelle, we also measured mitochondrial potential. Despite significant morphologic alterations, we found no effects of ACSVL3 KO on mitopotential. 97% of both U87MG and U87-KO cells were found to be alive, and less than 1% of these cells were depolarized. Further study is needed to clarify the relationship between ACSVL3 expression and mitochondrial structure and function.

Results presented here revealed that genetic KO of a relatively obscure enzyme of fatty acid metabolism, ACSVL3, produced far-reaching effects on cellular metabolism. Effects on lipid metabolism reported previously were anticipated, whereas changes in glucose metabolism and OXPHOS reported here were not. Our observations reinforce the notion that the integration and regulation of intermediary metabolism in U87MG and U87-KOcells is indeed complex, and worthy of further study.

## ACKNOWLEDGEMENTS

The authors thank Drs. G. William Wong and Xia Lei (Johns Hopkins University School of Medicine) for assistance with glucose uptake assays. The authors thank Drs. Akhilesh Pandey and Raja Sekhar Nirujogi, McKusick-Nathans Institute of Genetic Medicine, Johns Hopkins University School of Medicine for proteomic analysis of U87MG and U87-KO cells. Rabbit anti-Tom20 antibody was the kind gift of Dr. Hiromi Sesaki (JHUSOM). Supported by NIH 5R01NS062043 and Kennedy Krieger Institute institutional funds.

